# (*E*)-2-benzylidene-3-(cyclohexylamino)-2,3-dihydro-1*H*-inden-1-one (BCI) induces apoptosis via the intrinsic pathway in H1299 lung cancer cells

**DOI:** 10.1101/131045

**Authors:** Jong-Woon Shin, Sae-Bom Kwon, Yesol Bak, Sangku Lee, Do-Young Yoon

## Abstract

(*E*)-2-benzylidene-3-(cyclohexylamino)-2,3-dihydro-1*H*-inden-1-one (BCI) is known as a dual specific phosphatase 1/6 or MAPK inhibitor. However, its precise anti-lung cancer mechanism remains unknown. In this study, the effects of BCI on cell viability were investigated in the non-small cell lung cancer cell lines NCI-H1299, A549, and NCI-H460. We confirmed that BCI significantly inhibited the cell viability of NCI-H1299 compared to those of NCI-H460 and A549 cells. The anti-cancer effects of BCI were evaluated by MTS assay, annexin V-fluorescein isothiocyanate/propidium iodide staining, cell cycle analysis, reverse transcription-PCR, western blotting, and JC-1 staining in NCI-H1299 cells. BCI induced cellular morphological changes and inhibited viability of NCI-H1299 cells in a dose-dependent manner. BCI enhanced Bax expression and induced processing of caspase-9, caspase-3, and poly (ADP-ribose) polymerase as well as the release of cytochrome c from the mitochondria into the cytosol. BCI also down-regulated Bcl-2 expression but enhanced Bax expression in a dose-dependent manner in NCI-H1299 cells. In addition, BCI did not modulate death receptor expression or the extrinsic factor caspase-8 and Bid, a linker between the intrinsic and extrinsic apoptotic pathways in NCI-H1299 cells. On the basis of these results, we conclude that BCI induces apoptosis through a mediated intrinsic pathway, but not extrinsic pathway in NCI-H1299 cells. These results suggest that BCI can be used as a therapeutic agent in lung cancer.

## 1 Introduction

A large number of people die from lung cancer worldwide.^1^ Particularly, among patients with lung cancer, approximately 75% die from non-small cell lung cancer (NSCLC)^2^ and most are diagnosed at advanced, inoperable stages.^3^ Although surgery and chemotherapeutics can be employed, most patients with lung cancer eventually die. Therefore, new chemotherapeutic agents are needed to prolong the survival of patients with NSCLC.^4^

A tumor is a disease state characterized by uncontrolled cell proliferation and loss of apoptosis. Apoptosis is a genetically programmed, morphologically distinct form of cell death that can be triggered by a variety of physiological and pathological stimuli. There are distinct mechanisms that execute apoptosis according to various apoptotic stimuli. Apoptosis can be induced by two major pathways: the intrinsic pathway (mitochondria-dependent pathway) and extrinsic pathway (death receptor-dependent pathway).^4 5^ To maintain the balance between cell proliferation and death, homeostasis and apoptosis are programmed by our body system. The features of apoptosis are cell shrinkage and chromatin condensation, among others. The apoptotic process involves a cascade of events that inactivate critical survival pathways in multicellular organisms. Inhibition of apoptosis can prevent physiological cell death, which leads to the development and progression of tumor malignancy.^6 7^

Recent studies have revealed apoptosis as an ideal method for eliminating cancer cells.^8^ However, most cancer cells avoid apoptosis through genetic or morphological modifications. Defects in the apoptosis machinery provide a survival advantage to cancer cells and confer resistance of cancer cells to current anticancer therapies. Targeting critical apoptosis regulators is an attractive new cancer therapeutic strategy.^9^ Therefore, searching for agents that trigger apoptosis of tumor cells is a promising strategy in anti-cancer drug discovery.^10^

Many chemotherapeutic agents induce mitochondria-targeted apoptosis. The mitochondria-dependent (intrinsic) pathway is activated from inside the cell by severe cell stress, such as DNA or cytoskeletal damage, decreasing mitochondrial membrane potential, and transcription or post-translation activation of BH3-only proapoptotic B-cell leukemia/lymphoma 2 (Bcl-2) family proteins.^11^ The decreasing membrane potential induces the release of apoptotic proteins, including cytochrome c, from the mitochondria into the cytosol.^12^ Cytochrome c assembles with apoptotic protease-activating factor-1 (Apaf-1) to activate caspase 9. In turn, this caspase activates the effector caspases 3, 6, and 7 to carry out apoptosis.^11^

In this study, we assessed the effects of (*E*)-2-benzylidene-3-(cyclohexylamino)-2,3-dihydro-1*H*-inden-1-one (BCI), which is known as a dual specific phosphatase 1/6 or MAPK inhibitor and chemotherapeutic agent, on human lung cancer cells. We demonstrated the effects of BCI on apoptosis through the intrinsic pathway in NCI-H1299 cells.

## 2 Materials and methods

### 2.1 Reagents and antibodies

CellTiter 96 AQueous One Solution Cell Proliferation Assay Reagent [MTS, 3-(4, 5-dimethylthiazol-2-yl)-5-(3-carboxymethoxyphenyl)-2-(4-sulfophenyl)-2H-tetrazolium] was purchased from Promega (Madison, WI, USA). Propidium iodide (PI) was purchased from Sigma-Aldrich (St. Louis, MO, USA). NE-PER Nuclear and Cytoplasmic Extraction Reagents were purchased from Pierce (Rockford, IL, USA). Antibodies specific to PARP, caspase-3, caspase-8, caspase-9, p53, Bcl-2, Bcl-xL, Bax, Bid, pRb, p-pRb, and cytochrome C were purchased from Cell Signaling Technology (Beverly, MA, USA). The anti-rabbit IgG horseradish peroxidase (HRP)-conjugated secondary antibody and anti-mouse IgG HRP-conjugated secondary antibody were purchased from Millipore (Billerica, MA, USA). Antibodies specific to p27, p21, and glyceraldehyde 3-phosphate dehydrogenase (GAPDH) were purchased from Santa Cruz Biotechnology (Santa Cruz, CA, USA). JC-1 (5,50,6,60 -tetrachloro-1,10,3,30-tetraethyl benzimidazolycarbocyanine chloride) was purchased from Enzo (Farmingdale, NY, USA). General-caspase inhibitor Z-VAD-FMK was purchased from R&D Systems (Minneapolis, MN, USA). The FITC-Annexin V Apoptosis Detection Kit I was purchased from BD Biosciences (Franklin Lakes, NJ, USA).

### 2.2 Cell culture

Human NSCLC cell lines, including human lung adenocarcinoma cell line A549, human lung adenocarcinoma cell line NCI-H1299, and human large cell lung carcinoma cell line NCI-H460, were purchased from the American Type Culture Collection (Manassas, VA, USA). Cells were cultured in RPMI medium (Welgene Incorporation, Daegu, Korea) containing 10% (v/v) heat-inactivated fetal bovine serum (Hyclone, Logan, UT, USA). Cells were incubated at 37°C in an atmosphere of 5% CO_2_/95% air with saturated humidity.

### 2.3 Cell viability assays

Cell viability was assessed by the MTS dye reduction assay, which measures mitochondrial respiratory function. Lung cancer cells were seeded (1 × 10^4^ cells/mL) in 100 μL medium/well in 96-well plates, incubated overnight, and treated with various concentrations of BCI, as described in the figure legends, for 24 h. Cell viability was calculated by assessing MTS metabolism, as previously reported.^38^ Briefly, media samples (100 μL) were removed and incubated with 100 μL of MTS-PMS mix solution for 1 h at 37°C. Optical absorbance was measured at 492 nm using an ELISA reader (Apollo LB 9110, Berthold Technologies GmbH & Co. KG, Bad Wilbad, Germany).

### 2.4 Annexin V and propidium iodide staining

NCI-H1299 lung cancer cells (1.5 × 10^5^ cells/mL) were seeded into 60-mm culture dishes and incubated overnight. Cells were treated with BCI for 24 h, harvested using trypsin-EDTA, and washed with PBS. Annexin V and PI staining were performed using the FITC-Annexin V Apoptosis Detection Kit I according to the manufacturer’s instructions. Data were analyzed by flow cytometry, using a FACSCalibur instrument and CellQuest software (BD Biosciences).

### 2.5 Cell cycle analyses by flow cytometry

The cell cycle was analyzed by PI staining and flow cytometry. H1299 cells (1.5 × 10^5^ cells/mL) were seeded into 60-mm culture dishes and treated with various concentrations of BCI for 24 h. Cells were harvested with trypsin-EDTA and fixed with 80% ethanol. Next, the cells were washed twice with cold PBS and centrifuged, after which the resulting supernatants were discarded. The pellet was resuspended and stained with PBS containing 50 μg/mL PI and 100 μg/mL RNase A for 20 min in the dark. DNA content was analyzed by flow cytometry using a FACSCalibur instrument and CellQuest software.

### 2.6 Western blot analysis

Cells were treated with the specified concentrations of BCI for 24 h, harvested, washed with PBS, and recentrifuged (1,890 ×*g*, 5 min, 4°C). The resulting cell pellets were resuspended in lysis buffer containing 50 mM Tris (pH 7.4), 1.5 M sodium chloride, 1 mM EDTA, 1% NP-40, 0.25% sodium deoxycholate, 0.1% sodium dodecyl sulfate (SDS), and a protease inhibitor cocktail. The cell lysates were incubated on ice for 1 h and clarified by centrifugation at 17,010 ×*g* for 30 min at 4°C. Protein content was quantified by the Bradford assay (Bio-Rad, Hercules, CA, USA), using an UV spectrophotometer. Cell lysates were separated by 12–15% SDS polyacrylamide gel electrophoresis. Proteins were transferred onto polyvinylidene difluoride membranes (Millipore, Billerica, MA, USA), which were blocked in 5% non-fat dried milk dissolved in Tris-buffered saline containing Tween-20 (2.7 M NaCl, 53.65 mM KCl, 1 M Tris-HCl, pH 7.4, 0.1% Tween-20) for 1 h at room temperature. The membranes were incubated overnight at 4°C with specific primary antibodies. After washing, the membranes were incubated with the secondary antibodies (HRP-conjugated anti-rabbit or anti-mouse IgG) for 1 h at room temperature. After washing, the blots were analyzed using West-queen and a western blot detection system (iNtRON Biotechnology, SungNam, South Korea).

### 2.7 Nuclear and cytoplasmic fractionation

The BCI-treated cells were collected and fractionated using NE-PER Nuclear and Cytoplasmic Extraction Reagents (Thermo Fisher Scientific Inc., Waltham, MA, USA) according to the manufacturer’s protocol.

### 2.8 Analysis of mitochondrial membrane potential (MMP)

MMP (Δψm) was evaluated by JC-1 staining and flow cytometry. NCI-H1299 cells were seeded into 60-mm culture dishes (1.5 × 10^5^ cells/mL) and treated with various concentrations of BCI. Cells were harvested with trypsin-EDTA and transferred into 1.5-mL tubes. JC-1 (5 μg/mL) was added to the cells and mixed until it was completely dissolved, after which the cells were incubated in the dark for 10 min at 37°C in an incubator. The cells were centrifuged (300 ×*g*, 5 min, 4°C), washed twice with PBS, and resuspended in 200 μL PBS. The solutions were divided using a FACS Calibur instrument and analyzed by Cell Quest software. The protocol was performed in minimal light.

### 2.9 Reverse-transcription polymerase chain reaction (RT-PCR)

Cells treated with CTS were harvested. RNA was extracted using an easy-BLUETM Total RNA Extraction Kit (iNtRon Biotechnology, SungNam, Korea) according to the manufacturer’s instructions. cDNA products were obtained using M-MuLV reverse transcriptase (New England Biolabs, Ipswich, MA, USA). Each sample contained one of the following primer sets: FAS, 5′-AGG GAT TGG AAT TGA GGA AG-3′ (forward), 5′-ATG GGC TTT GTC TGT GTA CT-3′ (reverse); FASL, 5′-GCA GCC CTT GAA TTA CCC AT-3′ (forward), 5′-CAG AGG TTG GAC AGG GAA GAA-3′ (reverse); TRAIL, 5′-GTC TCT CTG TGT GGC TGT AA-3′ (forward), 5′-TGT TGC TTC TTC CTC TGG CT-3′ (reverse); GAPDH, 5′-GGC TGC TTT TAA CTC TGG TA-3′ (forward), 5′-TGG AAG ATG GTG ATG GGA TT-3′ (reverse); DR5, 5′-CAG AGG GAT GGT CAA GGT CG-3′ (forward), 5′-TGA TGA TGC CTG ATT CTT TGT GG-3′ (reverse); DR6, 5′-TGC AGT ATC CGG AAA AGC TC-3′ (forward), 5′-TCT GGG TTG GAG TCA TGG AT-3′ (reverse); DR3, 5′-CTA CTG CCA ACC ATG CCT AG-3′ (forward), 5′-TCG CCA TGT TCA TAG AAG CC-3′ (reverse); FADD, 5′-GGG GAA AGA TTG GAG AAG GC-3′ forward), 5′-CAG ATT CTC AGT GAC TCC CG-3′ (reverse); TRADD, 5′-CTA TTG CTG AAC CCC TGT CC-3′ (forward), 5′-AGA ATC CCC AAT GAT GCA CC-3′ (reverse);

### 2.10 Caspase inhibitor assay

The apoptosis mechanism was analyzed using caspase inhibitors. NCI-H1299 cells (1.5 × 10^5^cell/mL) were seeded into each well of a 60-mm cell culture dishes and treated with different 20 μM pan caspase inhibitors. After 3 h, 5 μM BCI was added to each plate. After 24 h, the cells were harvested and used in western blot analyses.

## 3 Results

### 3.1 BCI dose-dependently inhibits tumorigenic proliferation of H1299 lung cancer cells

The effect of BCI on the viability of several cell lines was determined by the MTS assay. We calculated the viability of BCI-treated cells compared to untreated control cells. Selected lung cancer cell lines (NCI-H1299, NCI-H460, and A549) were treated with various concentrations and for different time periods. As shown in Fig 1A, the viability of lung cancer cells was decreased in a dose-and time-dependent manner by BCI. The viability of NCI-H460 and NCI-H460 cells was slightly decreased by BCI (Fig 1). A549 cell viability was decreased by BCI for 24 h, but less than 48 h. The A549 cells were affected in a time-and dose-dependent manner, but these effects were greater in NCI-H1299 cells (Fig 1A–C). A549 and NCI-H460 cells were slightly inhibited by a high concentration of BCI (Fig 1B, C). Since BCI showed large effects on NCI-H1299 cells at 24 h of treatment, we used these cells in subsequent experiments. Thus, we examined the mechanism underlying apoptosis induced by BCI in NCI-H1299 lung cancer cells.

**Figure 1.**
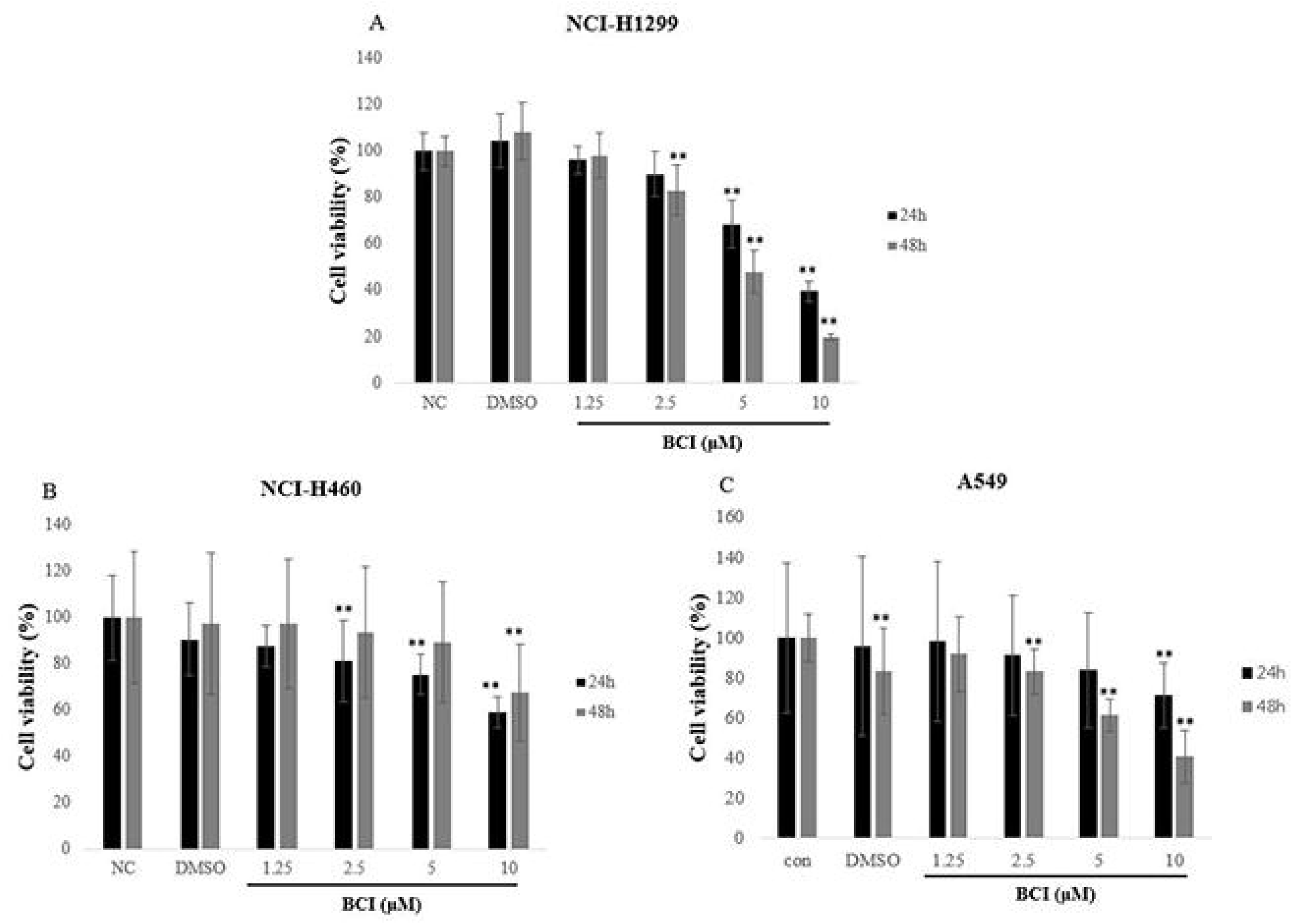
Effects of BCI on viability in human NSCLC cell lines. Cytotoxic effects of BCI on several lung cancer cell lines; (A) NCI-H1299, (B) NCI-H460, (C) A549 cells. These cells were treated for 24-48 h with various concentration of BCI. Cell viability was determined by MTS assay. The data are presented as the mean standard deviation. (n = 3) *p < 0.05, **p < 0.01 versus negative control groups.

### 3.2 BCI induces morphological changes and apoptosis in NCI-H1299 cells

Features of apoptosis include membrane shrinkage, nuclear fragmentation, and blebbing.^13^ Therefore, we evaluated changes to the cell membrane. Phase-contrast microscopy showed that BCI induced cell death and morphological changes in NCI-H1299 cells in a dose-dependent manner after a 24-h treatment (Fig 2A). When apoptosis is induced, the lipid phosphatidyl serine (PS) was translocated from the inner to outer cell membrane via a flip-flop movement. Annexin V, a calcium-dependent protein, binds PS with high affinity.^7^ Therefore, we performed annexin V-FITC/PI staining to evaluate changes in the orientation of PS. Annexin V-FITC/PI staining is generally used to detect apoptosis. NCI-H1299 cells treated with 1.25–5 μM BCI for 24 h showed a dramatically increased population of apoptotic cells compared to control cells, indicating that BCI induced apoptosis (Fig 2B). Thus, BCI may induce apoptosis in NCI-H1299 cells.

**Figure 2.**
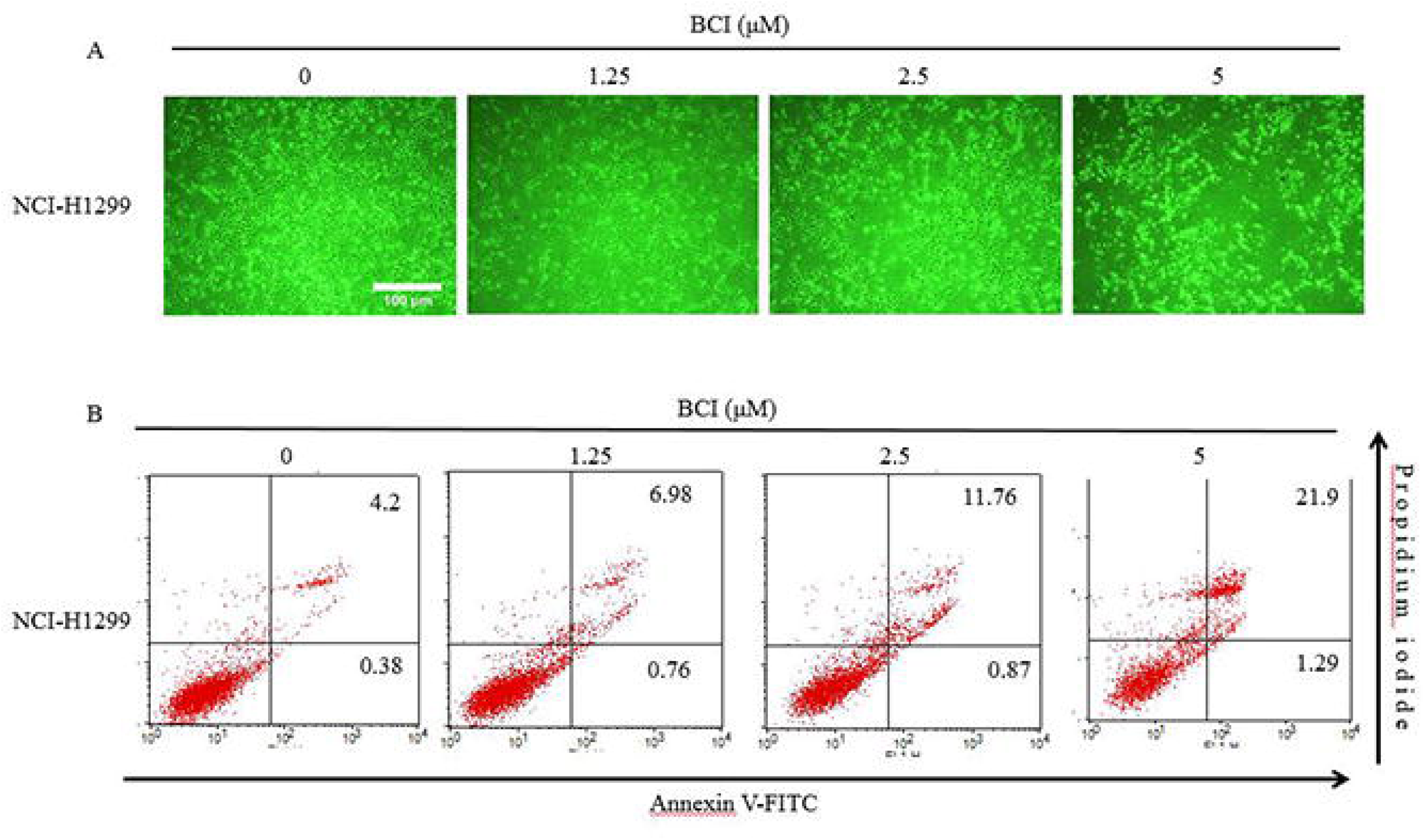
Effects of BCI on apoptosis in NCI-H1299 lung cancer cells. (A) Microscopic images of NCI-H1299 cells treated with BCI for 24 h. The photographs were taken by phase-contrast microscopy at a magnification of x100. BCI induces morphological changes in NCI-H1299 cells. (B) After treatment with the indicated concentration of BCI for 24 h, NCI-H1299 cells were stained with annexin V-FITC/PI.

### 3.3 BCI increases sub-G1 population

Next, we induced cell cycle progression to investigate the effects of BCI on cell cycle regulation. Cell cycle dysregulation is a major feature of cancer cells. Therefore, we examined whether BCI could regulate cell cycle in NCI-H1299 cells. To measure the effects of BCI on cell cycle progression, we assessed cell cycle progression by flow cytometry. The sub-G1 population contains hypodiploid fragmented DNA, which is a distinguishing feature of apoptosis. As shown in Fig 3A–B, compared to non-treated control cells, BCI-treated NCI-H1299 cells showed increased accumulation in subG1 phase. However, expression of cell cycle-related factors such as cyclin D1, cyclin E, and cyclin A were not significantly changed by BCI treatment in NCI-H1299 cells (Fig 3C). As shown in Fig 3D, p21 expression level was dose-dependently decreased by BCI. In contrast, p27 expression level was slightly increased by BCI. Since NCI-H1299 cells are p53 gene knock-out cells, p53 expression level was not detected by western blotting. In summary, these results show that BCI dramatically increased subG1 populations, but did not significantly increase cell proportion in the G0/G1, S, and G2 phases.

**Figure 3.**
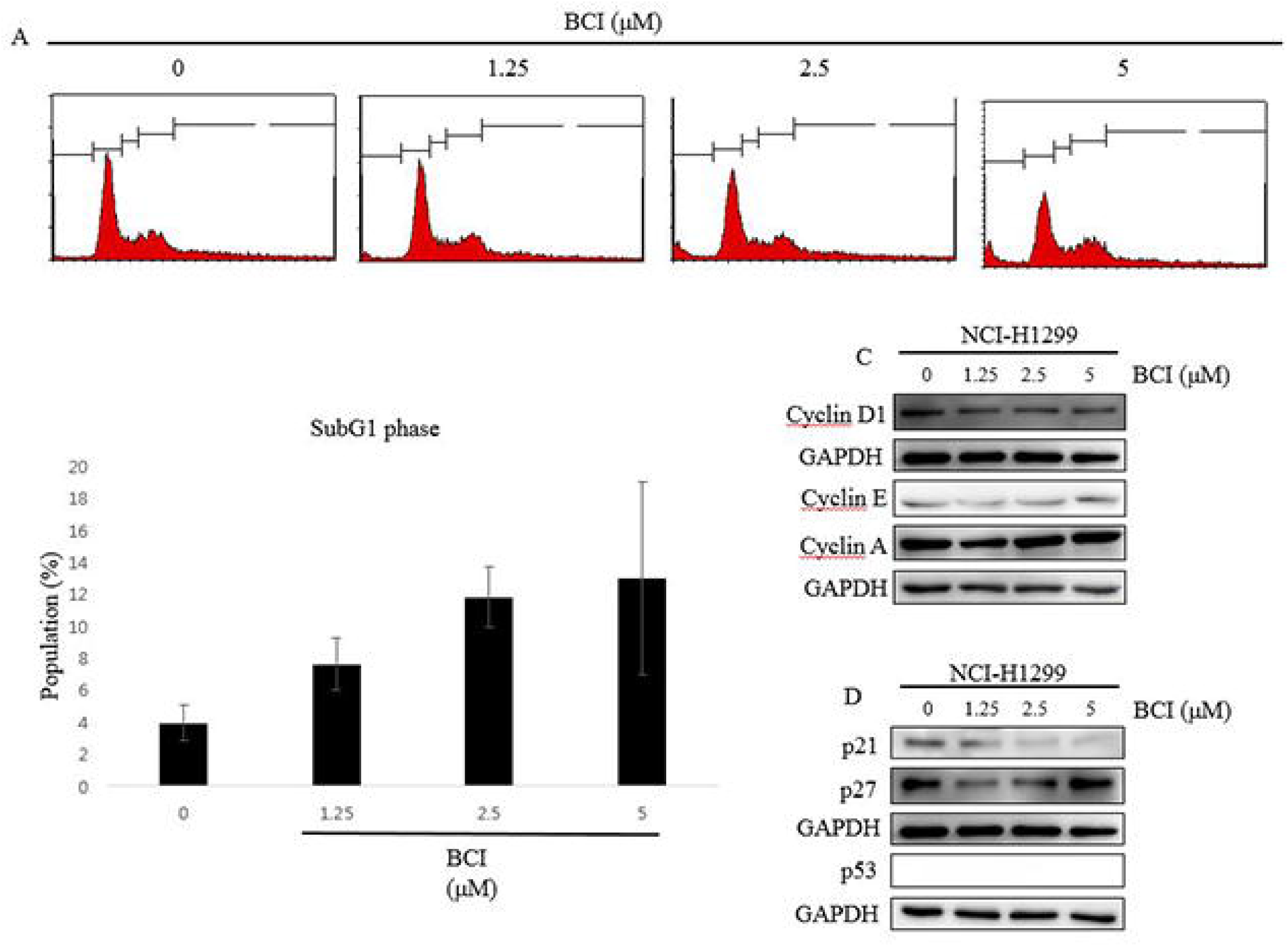
Effects of BCI on cell cycle progression in NCI-H1299 lung cancer cells. The cells were treated with various concentration of BCI in NCI-H1299 cells. After treatment with 0–5 μM BCI for 24 h, cells were fixed and stained with propidium iodine (PI). (A) Cell cycle profiles of BCI-treated NCI-H1299 cells. (B) The population of cells in various phases. (C) Western blots of cell cycle regulatory factors. GAPDH was used as an internal control.

### 3.4 BCI activates the intrinsic pathway by regulating the Bcl-2 family

Apoptosis is induced through two distinct signaling pathways; the death receptor pathway is typically triggered by ligation of death receptors such as Fas or tumor necrosis factor receptor, which recruits Fas-associated protein with death domain (FADD) and procaspase-8 to form a death-inducing signaling complex, leading to caspase-8 proteolytic activation. Activated caspase-8 can not only directly activate downstream caspase-3, but can also cleave Bid to truncated Bid (tBid), which in turn activates the mitochondrial pathway.^14^ Consistent with the results observed for caspase-8, BCI treatment did not affect the levels of DR3, DR5, DR6, FADD, TRAIL, TRADD, FasL, and Fas (Supplementary 1A). These data suggest that the extrinsic pathway is not involved in BCI-induced apoptosis. Next, we investigated whether the intrinsic pathway is involved in BCI-induced apoptosis. Mitochondrial dysfunction is the most important factor in the intrinsic pathway. The intrinsic pathway involves release of mitochondrial cytochrome C into the cytosol. Once released, cytochrome c combines with apoptotic protease activating factor 1 (Apaf-1) and procaspase-9 to form the apoptosome in the presence of ATP, resulting in activation of caspase-9 and caspase-3.^15^ To confirm that Bcl-2 family members play important roles in controlling the release of cytochrome c from the mitochondria, we performed western blotting in NCI-H1299 cells. Upon apoptotic signals, pro-apoptotic protein BAX was activated; in contrast, anti-apoptotic Bcl-2 was inhibited by BCI treatment in NCI-H1299 cells, but anti-apoptotic protein Bcl-xL was not affected (Fig 4C). Imbalance in the expression of pro-and anti-apoptotic proteins is associated with cell death.^16^ Since collapse of mitochondria membrane potential (MMP) is a characteristic feature of the intrinsic pathway, the effect of BCI on the MMP level was also examined by flow cytometry of JC-1-stained NCI-H1299 cells. JC-1 aggregation is a feature of healthy cells, whereas JC-1 monomers are a feature of apoptotic cells. As shown in Fig 4A, levels of the MMP in NCI-H1299 cells treated with various concentrations of BCI were significantly right-shifted compared to that in control cells. Thus, the levels of the MMP were decreased by BCI treatment in NCI-H1299 cells. To further confirm the release of mitochondrial cytochrome c into the cytosol, we evaluated cytochrome c by fractionation and western blotting in NCI-H1299 cells. As shown in Fig 4B, cytochrome c was released into the cytosol. Taken together, these results indicate that BCI induced apoptosis via the intrinsic pathway.

**Figure 4.**
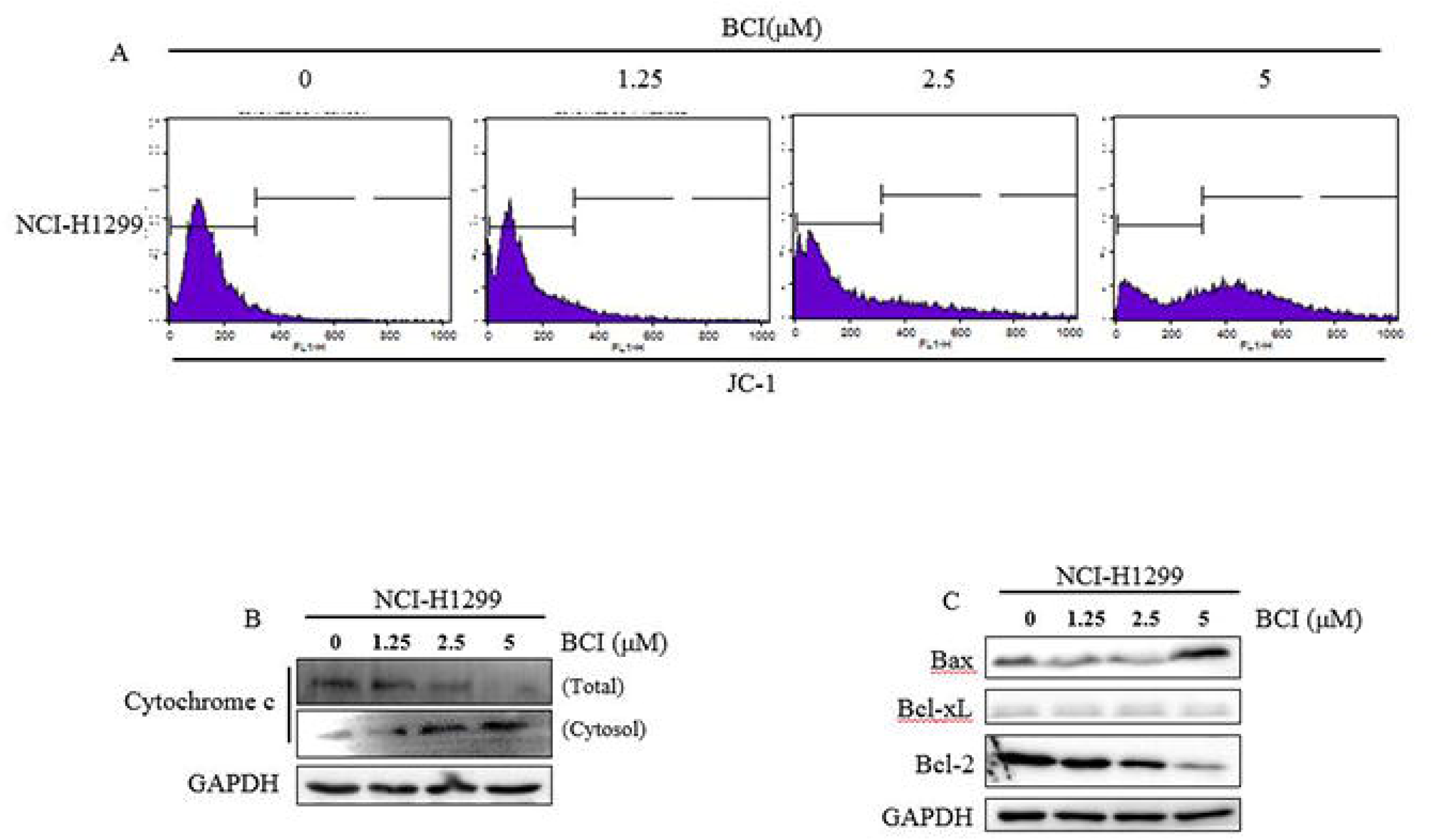
Effects of BCI on mitochondria membrane potential (MMP) in NCI-H1299 lung cancer cells. (A) Histogram profiles of JC-1 aggregation (FL-1, green) were obtained using flow cytometry. (B) Cytochrome C release was increased in a dose-dependent manner following BCI treatment. (C) Western blot analysis of the proapoptotic factors Bax and anti-apoptotic factors Bcl-xl and Bcl-2 in the NCI-H1299 cells following BCI treatment.

### 3.5 Effects of BCI on apoptosis related-factors in NCI-H1299 cells

To further investigate the apoptosis induced by BCI, we performed western blotting to evaluate apoptosis-related factors. A family of cysteine proteases known as caspases mediates apoptosis in mammalian cells.^17^ To maintain the apoptotic program under control, caspases were expressed in cells as inactive pro-caspases. At the time of apoptosis, initiator caspases such as caspase-9 and caspase-8 can be activated by cleavage. Furthermore, they cleave the precursor forms of effector caspases such as caspase-6, caspase-7, and caspase-3.^18^,^19^ In Fig 5A, caspase-9, which forms assembled apoptosome with Apaf-1 and cytochrome c, was cleaved by BCI in NCI-H1299 cells. Additionally, BCI treatment increased caspase-3 cleavage, followed by subsequent cleavage of poly (ADP-ribose) polymerase (PARP) (Fig 5A). Cleaved PARP is a major characteristic of programmed cell death.^17^ To confirm caspase-dependent apoptosis, we performed western blot analysis by Z-VAD-FMK, a pan caspase inhibitor. As shown in Fig 5B, pre-treatment with Z-VAD-FMK prevented BCI-induced cleavage of PARP and caspases such as caspase-3 and caspase-9. In summary, these results show that caspase-dependent apoptosis was induced by BCI.

**Figure 5.**
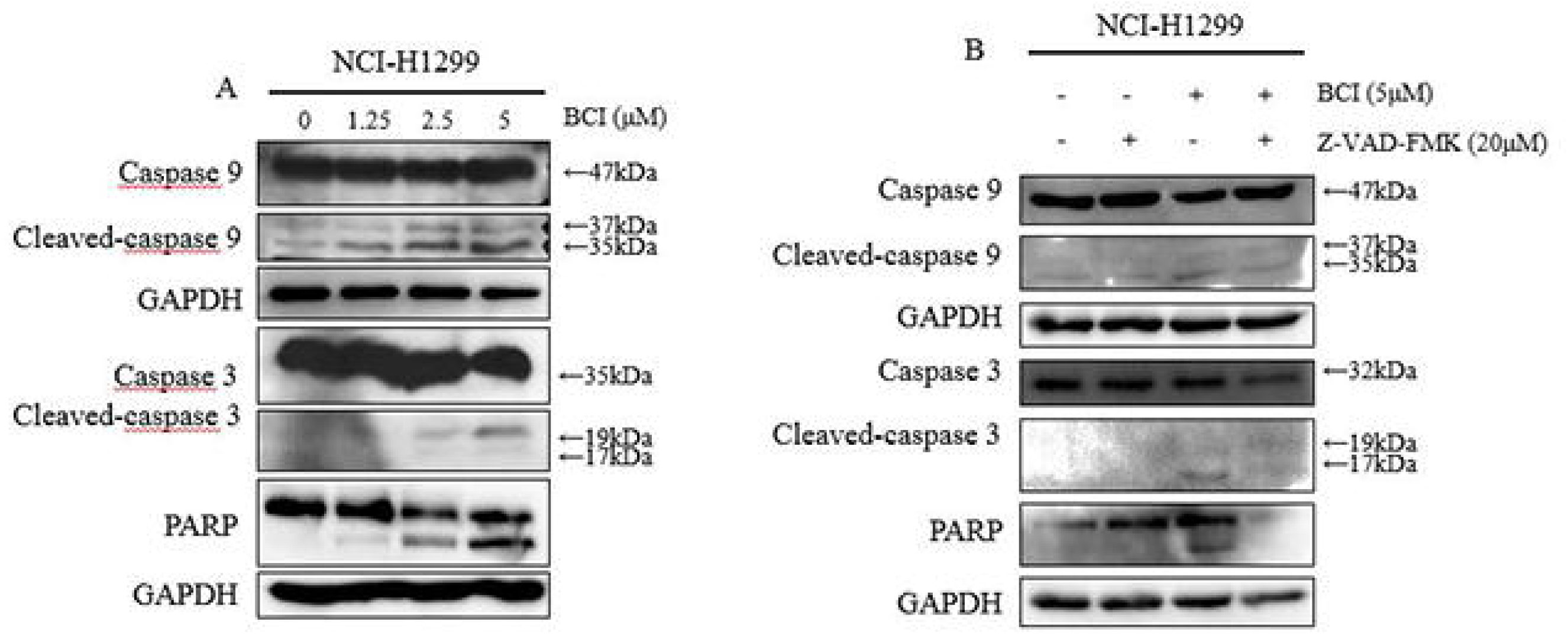
Effects of BCI on apoptosis related-factors in NCI-H1299 cells. NCI-H1299 cells were treated with BCI for 24 h. (A) Western blot of caspase 9, caspase 3, and poly (ADP-ribose) polymerase (PARP). GAPDH was used as an internal control. (B) Effects of caspase inhibitors on western blot, in BCI treated NCI-H1299 cells. The cells were pretreated with the pan-caspase inhibitor Z-VAD-FMK (20 μM) for 3 h. Next, the cells were treated with BCI for 24 h.

## 4 Discussion

BCI is a cell-permeable cylohexylamino-indenone compound that acts as an allosteric inhibitor against substrate binding-induced MAPK phosphatase activity of DUSP 1/6. Extracellular signal-regulated kinase phosphorylation is enhanced in HeLa cells overexpressing human Dusp 1 and Dusp 6. BCI also blocked Dusp 6 activity and enhanced fibroblast growth factor target gene expression in zebrafish embryos. *In vitro* studies supported a model in which BCI inhibits Dusp6 catalytic activation by ERK2 substrate binding. ^20^ However, it remains unclear how BCI induces apoptosis in lung cancer cells. Therefore, we evaluated the mechanism by which BCI induces apoptosis.

The main purpose of our study was to confirm the anti-cancer effect and associated mechanisms of BCI in human lung cancer cells. In this study, we investigated the anti-cancer effects of BCI in NCI-H1299 cells. We found that BCI inhibited tumor growth by inducing apoptosis in NCI-H1299 cells in a dose-dependent manner, as well as caspase activation and PARP cleavage (Fig 2A, Fig 5A–B). As shown in Fig 1A–C, the results showed greater anti-proliferative effect of BCI against NCI-H1299 cells than against NCI-H460 and A549 cells. Thus, NCI-H1299 cells were used as an experimental cell line because BCI was more effective in NCI-H1299 cells than in other cell lines. The present study evaluated the cytotoxic mechanism of BCI against NCI-H1299 cells.

Apoptosis can be induced by two major pathways: the intrinsic pathway or extrinsic pathway. The intrinsic pathway is induced by mitochondrial membrane permeabilization and cytochrome c release from the mitochondria into the cytosol.^21,22^ In the cytosol, cytochrome c was assembled into the apoptosome with Apaf-1 and caspase-9. Thus, activated-caspase-9 is an effector caspase.^23^ The extrinsic pathway receives signals through the binding of extracellular receptor ligand to proapoptotic death receptors located on the cell surface.^24^ Death receptors with ligand form a death-inducing signaling complex composed of death receptor, FADD, and caspase-8.^25^ These DISCs promote a downstream signaling cascade resulting in apoptosis.^26-28^ The imbalance between anti-apoptotic proteins such as Bcl-xl and Bcl-2 and pro-apoptotic proteins such as BAX and Bad determines the fate of cells.^29^ Mitochondria play an important role in death signal amplification and transduction of the apoptotic response. An important role of mitochondria in apoptotic signaling is the translocation of cytochrome c from the mitochondrial intermembrane into the cytosol. Once cytochrome c is released, cytochrome c together with Apaf-1 activate caspase-9, and the latter then activates caspase-3.^30^ The release of cytochrome c and cytochrome c-mediated apoptosis is regulated by Bcl-2 family members such as Bcl-2 and Bax, which are important regulators. In response to a variety of anticancer drugs, Bax translocates to the mitochondria and binds to the mitochondria membrane, allowing the release of cytochrome c.^31^ However, Bcl-2 prevents cytochrome c efflux by binding to the mitochondria membrane and forming a heterodimer with Bax, resulting in the neutralization of proapoptotic effects.^32^ Thus, the balance of Bcl-2 and Bax is important for determining cell fate, such as survival or death. We found that BCI treatment increased the ratio of Bax/Bcl-2. These results may be important in BCI-induced apoptosis.

In this study, we found that the process of BCI-induced apoptosis involved activation of caspase-3 and caspase-9, and treatment with a pan caspase inhibitor markedly prevented the BCI-induced cell apoptotic effects. Therefore, our results demonstrate that BCI-induced apoptosis depends on the activation of caspases, particularly caspase-3 and caspase-9 (Fig 5B). However, BCI treatment did not promote caspase-8 or its upstream molecules such as FasL and Fas. Taken together, BCI activates apoptosis through the intrinsic pathway, but not the extrinsic pathway. Apoptosis and cell cycle arrest are considered major cause of cell proliferation inhibition.^33^ When cells divide, the G2/M phase contains more DNA than the G0/G1 phase. The cells in SubG1 phase have hypodiploid fragmented DNA, which is an important marker of apoptosis.^34^ Cyclins such as cyclin D, A, and E promote cell cycle progression. Moreover, each cyclin is involved in a specific phase of the cell cycle.^35^ PI is an intercalating agent that cannot penetrate cell membranes of living cells, but can penetrate cell membranes of dead cells to distinguish living cells from dead cells.^36^ Our results suggest that BCI treatment did not induce cell cycle arrest in NCI-H1299 cells. However, BCI treatment increased sub-G1 populations in NCI-H1299 cells. To determine whether BCI induces cell death, we performed the annexin V-FITC/PI staining method. When apoptosis occurs, a flip-flop of lipid molecules occurs on the surface of the cell, and annexin V binds to the cell surface to detect cell death. ^37^ BCI-treated cells showed significantly induced apoptosis compared to untreated cells.

In conclusion, our results show that BCI has marked anti-cancer effects against lung cancer cells and increases subG1 populations in NCI-H1299 cells (Fig 3A–B). Our results revealed that BCI through the imbalance of Bcl-2 family proteins such as Bcl-2, Bax, and MMP, leads to apoptosis via the intrinsic pathway (Fig 6). This is the first study to demonstrate the molecular mechanism of BCI-mediated inhibition of lung cancer cell proliferation. Thus, BCI shows potential as a therapeutic agent for human lung cancer.

**Figure 6.**
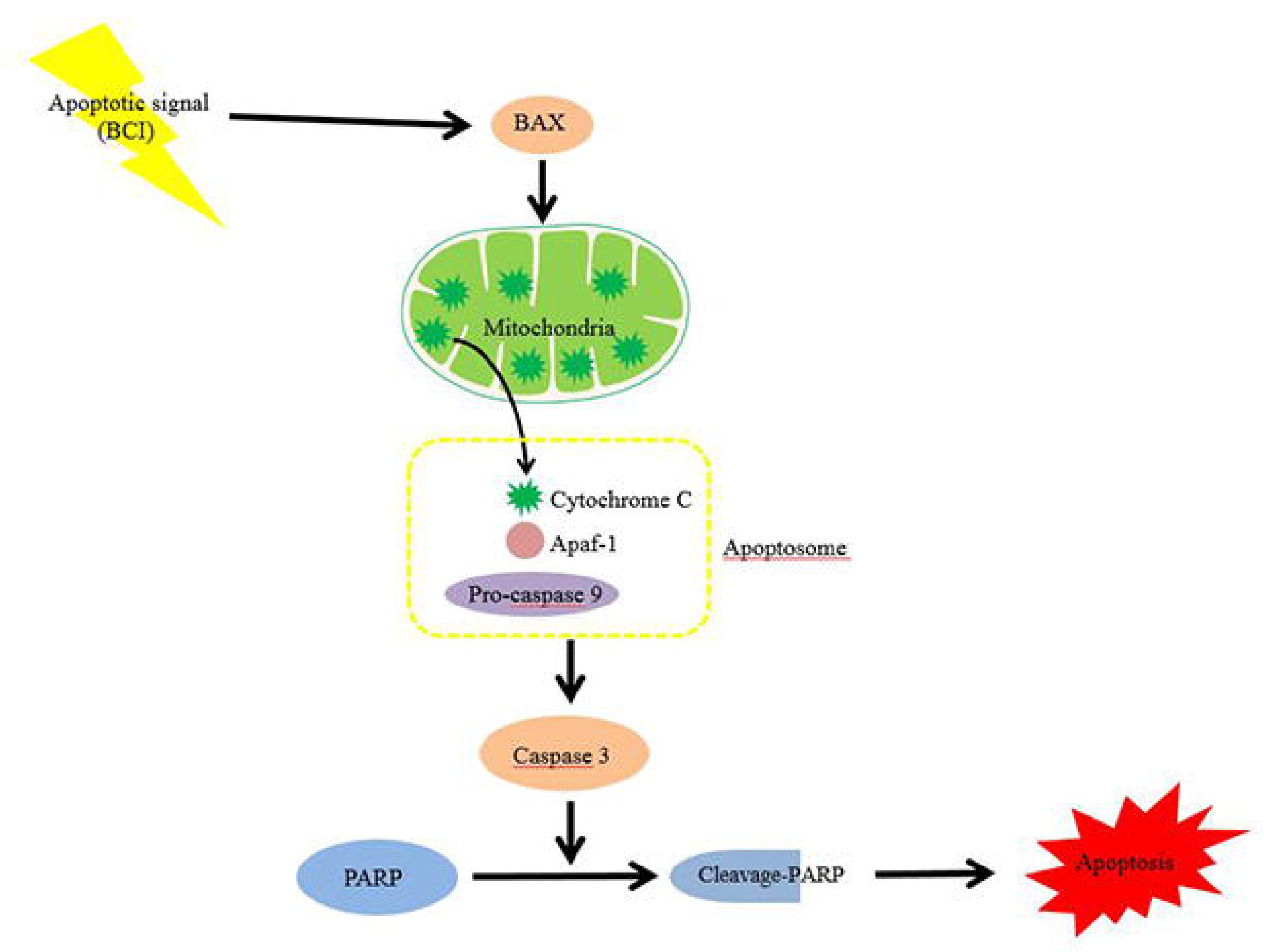
Schematic representation of potential BCI-induced apoptotic pathway in NCI-H1299 cells. The arrows symbolize activation of the signal pathways; the closed lines symbolize repression of the signals.

## ACKNOWLEDGEMENTS

This research was supported by Konkuk University in 2016.

## Reference

1 Siegel, R., Naishadham, D. & Jemal, A. Cancer statistics, 2013. CA Cancer J Clin 63, 11–30, doi:10.3322/caac.21166 (2013).

2 Jemal, A. et al. Global cancer statistics. CA Cancer J Clin 61, 69–90, doi:10.3322/caac.20107 (2011).

3 Pastorino, U. Lung cancer screening. Br J Cancer 102, 1681–1686, doi:10.1038/sj.bjc.6605660 (2010).

4 Ali, A. G., Mohamed, M. F., Abdelhamid, A. O. & Mohamed, M. S. A novel adamantane thiadiazole derivative induces mitochondria-mediated apoptosis in lung carcinoma cell line. Bioorg Med Chem 25, 241–253, doi:10.1016/j.bmc.2016.10.040 (2017).

5 Liu, D., Xiao, B., Han, F., Wang, E. & Shi, Y. Single-prolonged stress induces apoptosis in dorsal raphe nucleus in the rat model of posttraumatic stress disorder. BMC Psychiatry 12, 211, doi:10.1186/1471-244X-12-211 (2012).

6 Sipieter, F., Ladik, M., Vandenabeele, P. & Riquet, F. Shining light on cell death processes -a novel biosensor for necroptosis, a newly described cell death program. Biotechnol J 9, 224–240, doi:10.1002/biot.201300200 (2014).

7 Sankari, S. L., Masthan, K. M., Babu, N. A., Bhattacharjee, T. & Elumalai, M. Apoptosis in cancer--an update. Asian Pac J Cancer Prev 13, 4873–4878 (2012).

8 Jeong, S. Y. et al. Apoptosis induction of human leukemia cells by Streptomyces sp. SY-103 metabolites through activation of caspase-3 and inactivation of Akt. Int J Mol Med 25, 31–40 (2010).

9 Lu, J. et al. SM-164: a novel, bivalent Smac mimetic that induces apoptosis and tumor regression by concurrent removal of the blockade of cIAP-1/2 and XIAP. Cancer Res 68, 9384–9393, doi:10.1158/0008-5472.CAN-08-2655 (2008).

10 Xiao, J. X., Huang, G. Q., Zhu, C. P., Ren, D. D. & Zhang, S. H. Morphological study on apoptosis Hela cells induced by soyasaponins. Toxicol In Vitro 21, 820–826, doi:10.1016/j.tiv.2007.01.025 (2007).

11 Ashkenazi, A. Targeting the extrinsic apoptosis pathway in cancer. Cytokine Growth Factor Rev 19, 325–331, doi:10.1016/j.cytogfr.2008.04.001 (2008).

12 Fulda, S., Galluzzi, L. & Kroemer, G. Targeting mitochondria for cancer therapy. Nat Rev Drug Discov 9, 447–464, doi:10.1038/nrd3137 (2010).

13 Nagata, S. Apoptotic DNA fragmentation. Exp Cell Res 256, 12–18, doi:10.1006/excr.2000.4834 (2000).

14 Li, H., Zhu, H., Xu, C. J. & Yuan, J. Cleavage of BID by caspase 8 mediates the mitochondrial damage in the Fas pathway of apoptosis. Cell 94, 491–501 (1998).

15 Li, P. et al. Cytochrome c and dATP-dependent formation of Apaf-1/caspase-9 complex initiates an apoptotic protease cascade. Cell 91, 479–489 (1997).

16 Yang, W. L., Addona, T., Nair, D. G., Qi, L. & Ravikumar, T. S. Apoptosis induced by cryo-injury in human colorectal cancer cells is associated with mitochondrial dysfunction. Int J Cancer 103, 360–369, doi:10.1002/ijc.10822 (2003).

17 Alnemri, E. S. et al. Human ICE/CED-3 protease nomenclature. Cell 87, 171 (1996).

18 Salvesen, G. S. & Dixit, V. M. Caspases: intracellular signaling by proteolysis. Cell 91, 443–446 (1997).

19 Cryns, V. & Yuan, J. Proteases to die for. Genes Dev 12, 1551–1570 (1998).

20 Molina, G. et al. Zebrafish chemical screening reveals an inhibitor of Dusp6 that expands cardiac cell lineages. Nat Chem Biol 5, 680–687, doi:10.1038/nchembio.190 (2009).

21 Mignotte, B. & Vayssiere, J. L. Mitochondria and apoptosis. Eur J Biochem 252, 1–15 (1998).

22 Kroemer, G., Dallaporta, B. & Resche-Rigon, M. The mitochondrial death/life regulator in apoptosis and necrosis. Annu Rev Physiol 60, 619–642, doi:10.1146/annurev.physiol.60.1.619 (1998).

23 Rodriguez, J. & Lazebnik, Y. Caspase-9 and APAF-1 form an active holoenzyme. Genes Dev 13, 3179–3184 (1999).

24 Ashkenazi, A. & Dixit, V. M. Death receptors: signaling and modulation. Science 281, 1305–1308 (1998).

25 Kischkel, F. C. et al. Cytotoxicity-dependent APO-1 (Fas/CD95)-associated proteins form a death-inducing signaling complex (DISC) with the receptor. EMBO J 14, 5579–5588 (1995).

26 Algeciras-Schimnich, A. et al. Two CD95 tumor classes with different sensitivities to antitumor drugs. Proc Natl Acad Sci U S A 100, 11445–11450, doi:10.1073/pnas.2034995100 (2003).

27 Lavrik, I., Golks, A. & Krammer, P. H. Death receptor signaling. J Cell Sci 118, 265–267, doi:10.1242/jcs.01610 (2005).

28 Wilson, N. S., Dixit, V. & Ashkenazi, A. Death receptor signal transducers: nodes of coordination in immune signaling networks. Nat Immunol 10, 348–355, doi:10.1038/ni.1714 (2009).

29 van Gurp, M., Festjens, N., van Loo, G., Saelens, X. & Vandenabeele, P. Mitochondrial intermembrane proteins in cell death. Biochem Biophys Res Commun 304, 487–497 (2003).

30 Kroemer, G. & Reed, J. C. Mitochondrial control of cell death. Nat Med 6, 513–519, doi:10.1038/74994 (2000).

31 Li, Z. et al. Swainsonine activates mitochondria-mediated apoptotic pathway in human lung cancer A549 cells and retards the growth of lung cancer xenografts. Int J Biol Sci 8, 394–405, doi:10.7150/ijbs.3882 (2012).

32 Huang, Y. T., Huang, D. M., Chueh, S. C., Teng, C. M. & Guh, J. H. Alisol B acetate, a triterpene from Alismatis rhizoma, induces Bax nuclear translocation and apoptosis in human hormone-resistant prostate cancer PC-3 cells. Cancer Lett 231, 270–278, doi:10.1016/j.canlet.2005.02.011 (2006).

33 King, K. L. & Cidlowski, J. A. Cell cycle regulation and apoptosis. Annu Rev Physiol 60, 601–617, doi:10.1146/annurev.physiol.60.1.601 (1998).

34 Karna, P. et al. Polyphenol-rich sweet potato greens extract inhibits proliferation and induces apoptosis in prostate cancer cells in vitro and in vivo. Carcinogenesis 32, 1872–1880, doi:10.1093/carcin/bgr215 (2011).

35 Evans, T., Rosenthal, E. T., Youngblom, J., Distel, D. & Hunt, T. Cyclin: a protein specified by maternal mRNA in sea urchin eggs that is destroyed at each cleavage division. Cell 33, 389–396 (1983).

36 Moore, A., Donahue, C. J., Bauer, K. D. & Mather, J. P. Simultaneous measurement of cell cycle and apoptotic cell death. Methods Cell Biol 57, 265–278 (1998).

37 Vermes, I., Haanen, C., Steffens-Nakken, H. & Reutelingsperger, C. A novel assay for apoptosis. Flow cytometric detection of phosphatidylserine expression on early apoptotic cells using fluorescein labelled Annexin V. J Immunol Methods 184, 39–51 (1995).

38 Bak, Y. et al. A1E inhibits proliferation and induces apoptosis in NCI-H460 lung cancer cells via extrinsic and intrinsic pathways. Mol Biol Rep 40, 4507–4519, doi:10.1007/s11033-013-2544-0 (2013).

